# Evidence of multiple genome duplication events in *Mytilus* evolution

**DOI:** 10.1101/2021.08.17.456601

**Authors:** Ana Corrochano-Fraile, Andrew Davie, Stefano Carboni, Michaël Bekaert

## Abstract

Molluscs remain one significantly under-represented taxa amongst available genomic resources, despite being the second-largest animal phylum and the recent advances in genomes sequencing technologies and genome assembly techniques. With the present work, we want to contribute to the growing efforts by filling this gap, presenting a new high-quality reference genome for *Mytilus edulis* and investigating the evolutionary history within the Mytilidae family, in relation to other species in the class Bivalvia.

Here we present, for the first time, the discovery of multiple whole genome duplication events in the Mytilidae family and, more generally, in the class Bivalvia. In addition, the calculation of evolution rates for three species of the Mytilinae subfamily sheds new light onto the taxa evolution and highlights key orthologs of interest for the study of *Mytilus* species divergences.

The reference genome presented here will enable the correct identification of molecular markers for evolutionary, population genetics, and conservation studies. Mytilidae have the capability to become a model shellfish for climate change adaptation using genome-enabled systems biology and multi-disciplinary studies of interactions between abiotic stressors, pathogen attacks, and aquaculture practises.

## INTRODUCTION

The family Mytilidae constitutes a diverse group of bivalves, broadly distributed in marine environments. *Mytilus edulis* and *Mytilus galloprovincialis* are the common species cultivated in Europe and both hybridise with *Mytilus trossulus* where their geographical distribution overlaps (Gosling, 1992; Riginos and Cunningham, 2004) forming the European *Mytilus* Species Complex (Wilson et al., 2018). Nonetheless, a number of environmental and genetic barriers work together to maintain genetic discontinuities between the different species of the complex (El Ayari et al., 2019). *M. edulis* and *M. galloprovincialis* can be considered cosmopolitan species while *M. trossulus* remains more geographically confined to the northernmost regions of the Pacific and Atlantic oceans and to the Baltic Sea (Gosling, 1994). At a finer geographical scale, mussel species display an extraordinary capability of environmental adaptation, extending from high inter-tidal to sub-tidal regions, from estuary to fully marine conditions, and from sheltered to extremely wave-exposed shores. Mussels are furthermore exposed to a wide range of potentially pathogenic microorganisms and pollutants, and yet they display a remarkable resilience to stress and infections (Gerdol et al., 2020). Of particular interest are observations of a relatively high heterozygosity, rapid evolutionary responses to environmental threats, including predation (Freeman, 2006), and recent suggestions that widespread relaxed selection in “low locomotion” molluscs, such as bivalves, and high copy number variants (Modak et al., 2021) could underpin observed high resilience and rapid adaptation to new environments (Sun et al., 2017).

The Phylum Mollusca remains significantly under-represented amongst those with available genomic resources (Sigwart et al., 2021). To date, only two high-quality genomes and associated gene models are available within the Mytilidae family: *M. galloprovincialis*, first sequenced by Murgarella et al. (2016) and then improved by Gerdol et al. (2020), and *M. coruscus* recently sequenced by Li et al. (2020) and (Yang et al., 2021). Comparative genomics provides an opportunity to investigate the “signatures” that natural selection has left on the genomes of related species. By analysing the frequency distribution of synonymous substitution per synonymous sites, it is possible to identify major evolutionary events, including Whole-Genome Duplications (WGDs). While WGDs are rare within animal lineages, they deeply shaped vertebrate evolution and represent important evolutionary landmarks from which some major lineages have diversified (Berthelot et al., 2014). Furthermore, comparisons among related species adapted to contrasting niches, can provide an opportunity to investigate how their genomes diverge in response to different habitat conditions (Oliver et al., 2010). In cases where specific amino acids are known to affect protein function, analyses of intra-specific polymorphism and divergence can be used to directly study function variation in natural populations (Dean and Thornton, 2007; Storz et al., 2015). Both whole genome duplication analysis and positive selection genome-wide analysis can therefore expose the strength and direction of selection on genes’ functional variation and corroborate the adaptive significance of the loci under study (Linnen et al., 2009; Natarajan et al., 2015).

With this work, we want to contribute to the growing efforts in filling the gap in Molluscs genomic resources (Davison and Neiman, 2021) by presenting a new high-quality reference genome for *M. edulis* and investigating the evolutionary history within the Mytilidae family and in relation to other species in the class Bivalvia. Here we present a new reference genome for *M. edulis*; we introduce the first evidence of WGD events in the Mytilidae family and in Bivalvia more generally; finally, we identify gene clusters under significant positive selection within each species of the Mytilidae family for which suitable reference genomes are available (*M. edulis, M. galloprovincialis* and *M. coruscus*).

The availability of this reference genome will not only increase interest in Mytilidae as a model for ecological and evolutionary research, but will be also a valuable tool for breeding programmes (Regan et al., 2021). The discovery of multiple duplication events will enable the correct identification of molecular markers for evolutionary, population genetics, and conservation studies. The Mytilidae have the capability to become a model shellfish for climate change adaptation using genome-enabled systems biology and multi-disciplinary studies of interactions between abiotic stressors, pathogen attacks, and aquaculture practises.

## MATERIALS AND METHODS

### Material collection

The *M. edulis* used in this work was obtained from a female adult blue mussel gill tissue from a location previously reported to present a pure *M. edulis* population (Wilson et al., 2018). Gill tissue was dissected, stored in 95% ethanol and shipped to Novogene Ltd (Cambridge, UK) for DNA extraction and sequencing.

### Library construction and sequencing

High-quality DNA was used for subsequent library preparation and sequencing using both the PromethION and Illumina platforms at Novogen UK (Novogene UK Company Ltd, UK). To obtain long non-fragmented sequence reads, 15 µg of genomic DNA was sheared and size-selected (30-80 kb) with a BluePippin and a 0.50% agarose Gel cassette (Sage Science, USA). The selected fragments were processed using the Ligation Sequencing 1D Kit (Oxford Nanopore, UK) as directed by the manufacturer’s instructions and sequenced using the PromethION DNA sequencer (Oxford Nanopore, UK) for 48 hours. For the estimation and correction of genome assembly, an Illumina DNA paired-end library with an insert size of 350 bp was built in compliance with the manufacturer’s protocol and sequenced on an Illumina HiSeq X Ten platform (Illumina Inc., USA) with paired-end 150 nt read layout.

### RNA isolation, cDNA library construction and sequencing

The total RNA was extracted using the TRIzol reagent (Invitrogen, USA) according to the manufacturer’s instructions. The preparation and sequencing reactions of cDNA library were done by Novogene Ltd. Briefly, the poly (A) messenger RNA was isolated from the total RNA with oligo (dT) attached magnetic beads (Illumina Inc., USA). Fragmentation was carried out using divalent cations under elevated temperature in Illumina proprietary fragmentation buffer. Double-stranded cDNAs were synthesised and sequencing adaptors were ligated according to the Illumina manufacturer’s protocol (Illumina Inc., USA). After purification with AMPureXP beads, the ligated products were amplified to generate high quality cDNA libraries. The cDNA libraries were sequenced on an Illumina Hiseq 4000 platform (Illumina Inc., USA) with paired-end reads of 150 nucleotides.

### Genome assembly

Reads from the two types of sequencing libraries were used independently during assembly stages. Long-reads were filtered for length (> 5,000 nt) and complexity (entropy over 15), while all short reads were filtered for quality (QC > 25), length (150 nt), absence of primers / adaptors and complexity (entropy over 15) using fastp v0.20.1 (Chen et al., 2018). Using Jellyfish v2.3.0 (Marçais and Kingsford, 2011), the frequency of 17-mers and 23-mers in the Illumina filtered data was calculated with a 1 bp sliding window (Vurture et al., 2017) to evaluate genome size. Long-reads were then assembled using wtdbg2 v2.5 (Ruan and Li, 2020) which uses fuzzy Bruijn graph as well as raven v1.5.0 (Vaser and Šikić, 2021). As it assembles raw reads without error correction and then creates a consensus from the intermediate assembly outputs, several error corrections, gap closing, and polishing steps have been implemented. The initial outputs have been combined using Quickmerge v0.3 (Solares et al., 2018). The combined output was re-aligned to the long-read and polished using Minimap2 v2.17 (Li, 2018) and Racon v1.4.3 (Vaser et al., 2017), first with filtered reads, to bridge potential gaps, then with the filtered reads to correct for error. Finally, Pilon v1.23 (Walker et al., 2014) was used to polish and correct for sequencing error using the short-reads. The redundant contigs due to diploidy were reduced by aligning the long reads back to the assembly with Minimap2 v2.17 (Li, 2018) and by passing the alignment through the Purge Haplotigs pipeline v1.1.1 (Roach et al., 2018). This reduced the artefact scaffolds and created the final haploid representation of the genome. Scaffolds were ordered with Medusa v1.6 (Bosi et al., 2015) using *M. galloprovincialis* (Gerdol et al., 2020; Murgarella et al., 2016) and *M. coruscus* (Li et al., 2020) genome scaffolds. Mitochondrial genome was annotated using MITOS r999 (Bernt et al., 2013) and manually curated.

### Repeat sequences

The transposable elements were annotated using a de novo prediction using RepeatModeler v2.0.1 (Flynn et al., 2020) and LTR-Finder v1.07 (Stanke et al., 2008). The repetitive sequences yielded from these two programs were combined into a non-redundant repeat sequence library. With this library, the *M. edulis* genome was scanned using RepeatMasker v4.10 (Smit et al., 2019) for the representative sequences.

### Gene models

RNA-seq reads of poor quality (*i*.*e*. with an average quality score less than 20) or displaying ambiguous bases or too short and PCR duplicates were discarded using fastp v0.20.1 (Chen et al., 2018). Ribosomal RNA was further removed using SortMeRNA v3.0.2 (Kopylova et al., 2012) against the Silva version 119 rRNA databases (Quast et al., 2012). The cleaned RNA-seq reads were pooled and mapped to the genome using the using HiSat2 v2.2.0 (Kim et al., 2019). A combined approach that integrates ab initio gene prediction and RNA-seq-based prediction was used to annotate the protein-coding genes in *M. edulis* genome. We used Braker v2.1.5 (Hoff et al., 2019) to make *de novo* gene predictions. The accuracy and sensitivity of the predicted model was improved by applying iterative self-training with transcripts. The predicted coding sequences were been annotated using the InterProscan v5.46-81.0 (Jones et al., 2014; Mitchell et al., 2019), Swiss-Prot release 2020 02 (Bateman et al., 2017) and Pfam release 33.1 El-Gebali et al. (2019) databases. For classification, the transcripts were handled as queries using Blast+/BlastP v2.10.0 (Altschul et al., 1990), E-value threshold of 10^−5^, against Kyoto Encyclopedia of Genes and Genomes (KEGG) r94.1 (Kanehisa et al., 2019). Gene Ontology (Ashburner et al., 2000) was recovered from the annotations of InterPro, KEGG and SwissProt. Subsequently, the classification was performed using R v4.0.0 (R Core Team, 2021) and the Venn diagram was produced by jvenn (Bardou et al., 2014). The completeness of gene regions was further tested using BUSCO v4.0.2 (Simão et al., 2015) with a Metazoa (release 10) benchmark of 954 conserved Metazoa genes.

### Phylogenetic Tree

Concatenated alignments constructed from all mitochondrial shared CDS sequences were used to construct a phylogenetic tree. Sequences were aligned using MACSE v10.02 (Ranwez et al., 2011). A Maximum Likelihood (ML) tree was inferred under the GTR model with gamma-distributed rate variation (Γ) and a proportion of invariable sites (I) using a relaxed (uncorrelated log-normal) molecular clock in RAxML v8.2.12 (Stamatakis, 2014).

### Calculating Ka, Ks, and Ka/Ks values

Complete annotated nuclear genomes of Bivalvia (class) were collected. Blast+/BlastP v2.10.0 (Altschul et al., 1990) was used to search for duplicated sequences in protein-coding genes between each genome. Duplicate pairs were identified as sequences that demonstrated over 70% sequence similarity, mutual protein coverage > 80%, protein length > 30 amino-acid from an all-against-all search. Duplicated sequences were aligned accounting for codon and coding frame, using MACSE v10.02 (Ranwez et al., 2011). Finally, the Ka (number of non-synonymous substitutions per non-synonymous site) and Ks (number of synonymous substitutions per synonymous site) for each pair was calculated using an MPI version of KaKs Calculator (Zhang et al., 2006) under the MLPB model (Tzeng, 2004). The Ks values > 5.0 were excluded from further analysis due to the saturated substitutions at synonymous sites. Univariate mixture models were fitted to the distributions of Ks by expectation maximisation that uses the finite mixture expectation maximisation algorithm (Benaglia et al., 2009; Tiley et al., 2018).

### Estimation of evolution rates

A set of core-orthologs was constructed from the three complete annotated nuclear genomes of Mytilinae (subfamily), *M. edulis, M. galloprovincialis* (Gerdol et al., 2020) and *M. coruscus* Li et al. (2020) and were used to identity cluster of orthologous genes with a 1:1:1 ratio. Ka and Ks estimation were reported pairwise between species after MACSE v10.02 (Ranwez et al., 2011) and KaKs Calculator (Zhang et al., 2006) under the MLPB model (Tzeng, 2004) as above. From a literature search of comparative mussel biology studies, we identified genes relevant to core physiological functions, specifically immunity, stress response and shell formation. Subsequently, a local BLAST search was conducted on NCBI to identify the genes of interest in the available genomes of the three species with a cut-off point of 80% sequences similarity.

## RESULTS

### Sequencing results

After sequencing with the PromethION platform, a total of 15.95 million (111.65 Gb) long-reads were generated and used for the genome assembly. The mean length of the sequences was 7,002 nt. The Illumina HiSeq X Ten platform produced 652.47 million (195.74 Gb) paired-ended short reads (150 nt). Based on the presumption that the genome size will be similar to that of closely related taxa: *M. galloprovincialis* (Murgarella et al., 2016) and *M. coruscus* (Li et al., 2020) with an estimated genome size of 1.60 Gb and 1.90 Gb respectively; therefore, the estimated sequencing coverage was 64x and 113x, for long and short reads respectively (Table S1).

### De novo assembly of the M. edulis genome

Using Jellyfish, the frequency of 17-mers and 23-mers in the Illumina filtered data were determined (Fig. S1) and followed the theoretical Poisson distribution typical of a diploid species (Benadelmouna et al., 2018). The proportion of heterozygosity in the *M. edulis* genome was evaluated as being 3.69% and 4.84% respectively, and the genome size was estimated as 1.01 Gb and 1.10 Gb, with a repeat content of 68.13% and 39.91% respectively (Table 1).

**Table 1.** Statistics of the genome assembly of *M. edulis*.

| Category | Number/length |
| --- | --- |
| K-mer = 17 |  |
| Estimated genome size | 1,010,184,781 nt |
| Estimated repeats | 688,190,885 nt |
| Estimated heterozygosity | 3.69% |
| K-mer = 23 |  |
| Estimated genome size | 1,096,306,163 nt |
| Estimated repeats | 437,569,400 nt |
| Estimated heterozygosity | 4.84% |
| Number of contigs | 3,339 |
| Total length | 1,827,085,763 nt |
| Total repeats | 1,029,206,554 nt |
| Observed heterozygosity | 0.48% |
| Largest contig | 10,529,124 nt |
| N<sub>50</sub> | 1,097,279 nt |
| GC | 32.17% |
| Read Mapped | 91.35% |
| Avg. coverage depth | 152x |
| Coverage over 10x | 99.99% |
| N's per 100 kbp | 13.73 |
| BUSCO recovered | 98.9% |
| Predicted rRNA genes | 132 |
| Predicted protein coding genes | 69,246 |

Long-reads were assembled, polished with Racon and short-read sequence were corrected with Pilon, creating an assembled genome of 3,339 contigs with a total length and contig N_50_ of 1.83 Gb and 1.10 Mb, respectively (Table 1). The realignment of the short-reads also provided descriptions of the mean observed heterozygosity of 0.48%, which is consistent with the most recent evidence collected from de novo RAD analysis (Vendrami et al., 2020) and microsatellite loci study (Coolen et al., 2020).

### Repeat sequences and Gene models

The transposable elements and repetitive sequences have been annotated using RepeatMasker and LTR-Finder. In total, we have found 1.03 Gb (56.33%) of repetitive sequences (Table S2). We used a combined method that integrates *ab initio* gene prediction and RNA-seq-based prediction to annotate the protein-coding genes in *M. edulis* genome. A total of 69,246 distinct gene models and 73,842 transcripts were annotated. The completeness of gene regions was further assessed using BUSCO using a Metazoa (release 10) benchmark of 954 conserved Metazoa genes, of which 93.8% had complete gene coverage (including 29.4% duplicated ones), 5.1% were fragmented and only 1.1% were classified as missing (Fig. 1A). These data largely support a high-quality *M. edulis* genome assembly, which can be used for further investigation.

**Figure 1.**
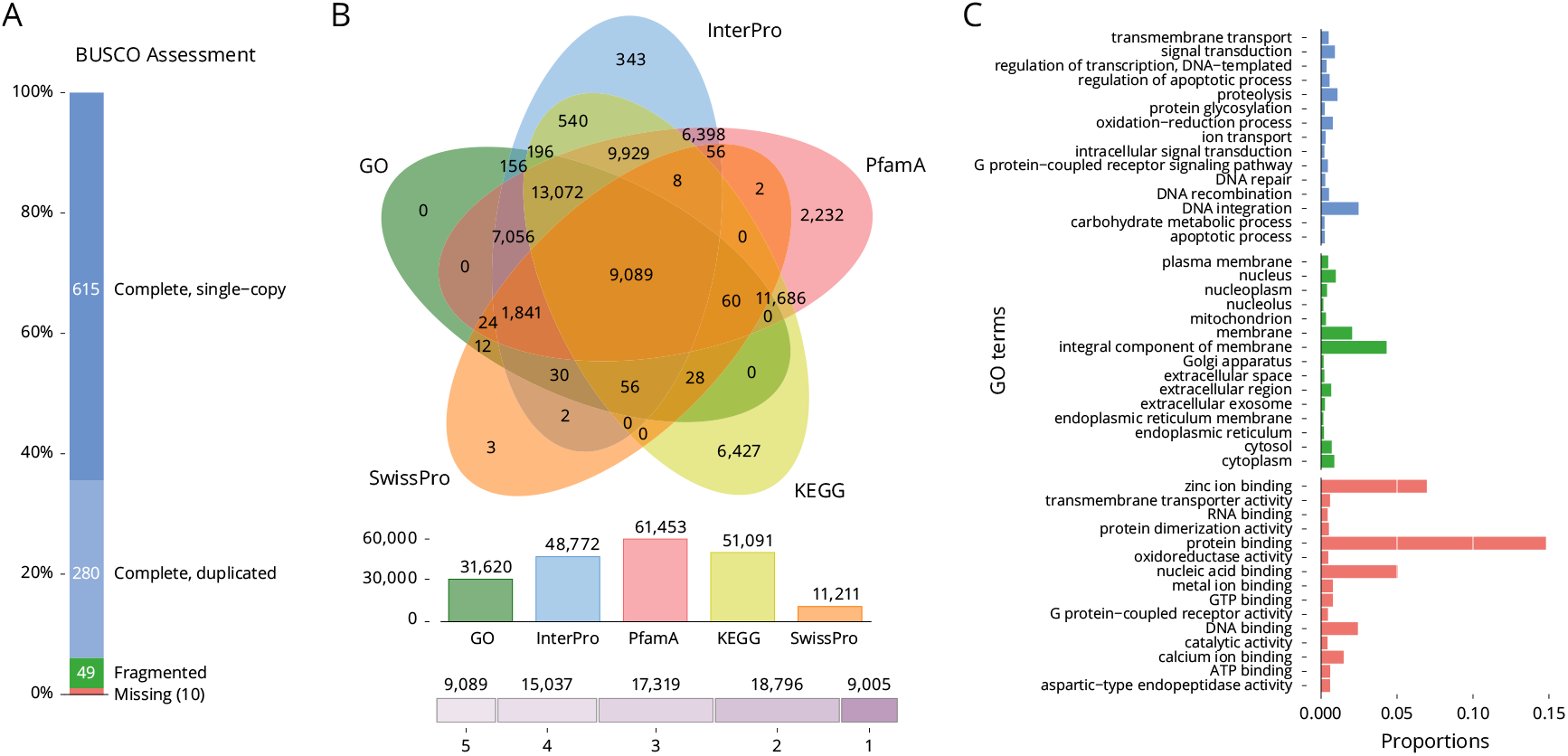
Gene composition and annotation estimations. **A** BUSCO evaluation (Metazoa database; number of framework genes 954), 98.9% of the gene were recovered; **B** A five-way Venn diagram. The figure shows the unique and common genes displaying predicted protein sequence similarity with one or more databases (details in Supplementary Table S3); **C** Level 2 GO annotations using the gene ontology of assembled transcripts.

The predicted proteins from the reconstructed genes were subjected to BlastP similarity searches against SwissProt, Pfam, InterPro, KEGG and GO databases. Of the total 69,246 gene models annotated by at least one database, 9,005 (13.0%) were annotated in all five databases used (Fig. 1B and Table S3). A total of 31,620 predicted genes were annotated to three major GO classes: “biological processes”, “cellular components” and “molecular functions” (Fig. 1C).

### Mitochondrial genome

The mitochondrial genome was retrieved manually from the genome assembly. The sequence of 16.74 kb was validated for continuity and circularity, and fully annotated. The full mitochondrial genome (Fig. 2A) was compared to the reference *M. edulis* genome (Boore et al., 2004). Only one haplotype was recovered which is identical at 99% (85 SNPs) with the reference genomes (EBI Accession NC 006161.1) and is consistent with a female mitotype (Breton et al., 2006). Complete annotated mitochondrial genomes for all Mytilinae (subfamily) to date (11 species; Table S4) were collected. Concatenated alignments constructed from all mitochondrial shared CDS sequences were used to construct a phylogenetic tree (Fig. 2B). This phylogenetic tree is consistent with the species relationships observed in previous studies (Lee et al., 2019).

**Figure 2.**
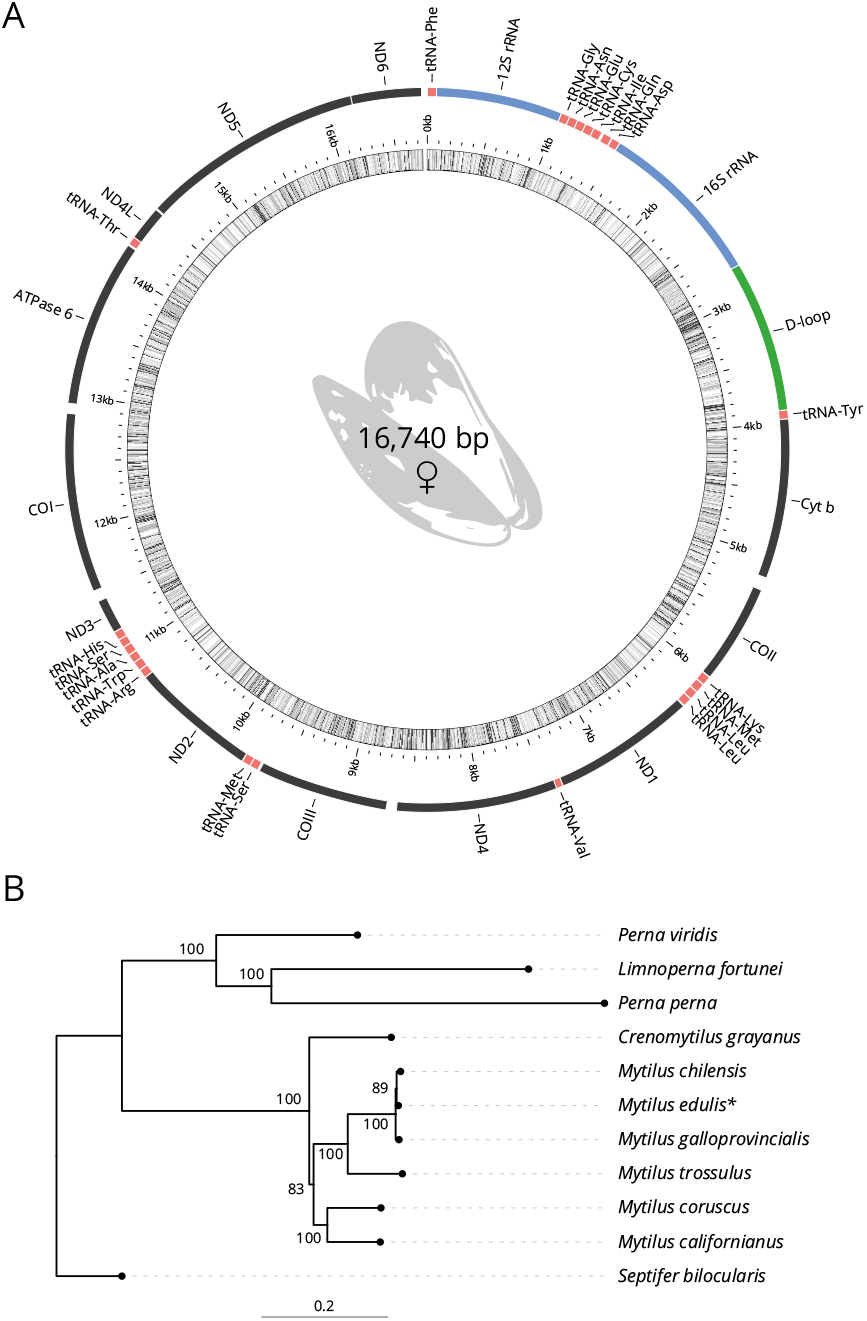
Genome assembly. **A** *M. edulis* annotated mitochondrial genome; **B** Phylogenetic tree inferred from the mitochondrial gene.

### Detecting whole genome duplications

To assess the paleo-history of the Mytilinae (subfamily), we performed a comparative genomic investigation. A total of 2,293 gene duplications younger than Ks = 5 were inferred across the total data set of 16,291 assembled unigene clusters in Mytilinae (*M. coruscus, M. edulis* and *M. galloprovincialis*). The histograms of duplication ages for each species analysed demonstrated evidence of two large-scale duplications (Fig. 3A). Mixture model analyse of Ks distributions (Fig. 3A and Table S5) to identify ancient whole genome duplications (Shi et al., 2010; Tiley et al., 2018) were consistent with the two consecutive whole genome duplication events (*α*-WGD and *β* -WGD). The duplication distributions each contained evidence of two peaks of similar synonymous divergences. For example, in *M. edulis* these peaks are located at median Ks of 0.6132 and 1.8196 (Table S5). We extended the analysis to all Bivalvia (class). Out of the 45 whole genomes available, only 7 (including the three Mytilinae) have gene models allowing further analyses (Table S6). All exhibit evidence of *α*-WGD and *β* -WGD (Table S5). The median value, for Ks peaks, is compatible with a shared WGD event (compatible age) indicating that these taxa diverged after their most recent WGDs.

**Figure 3.**
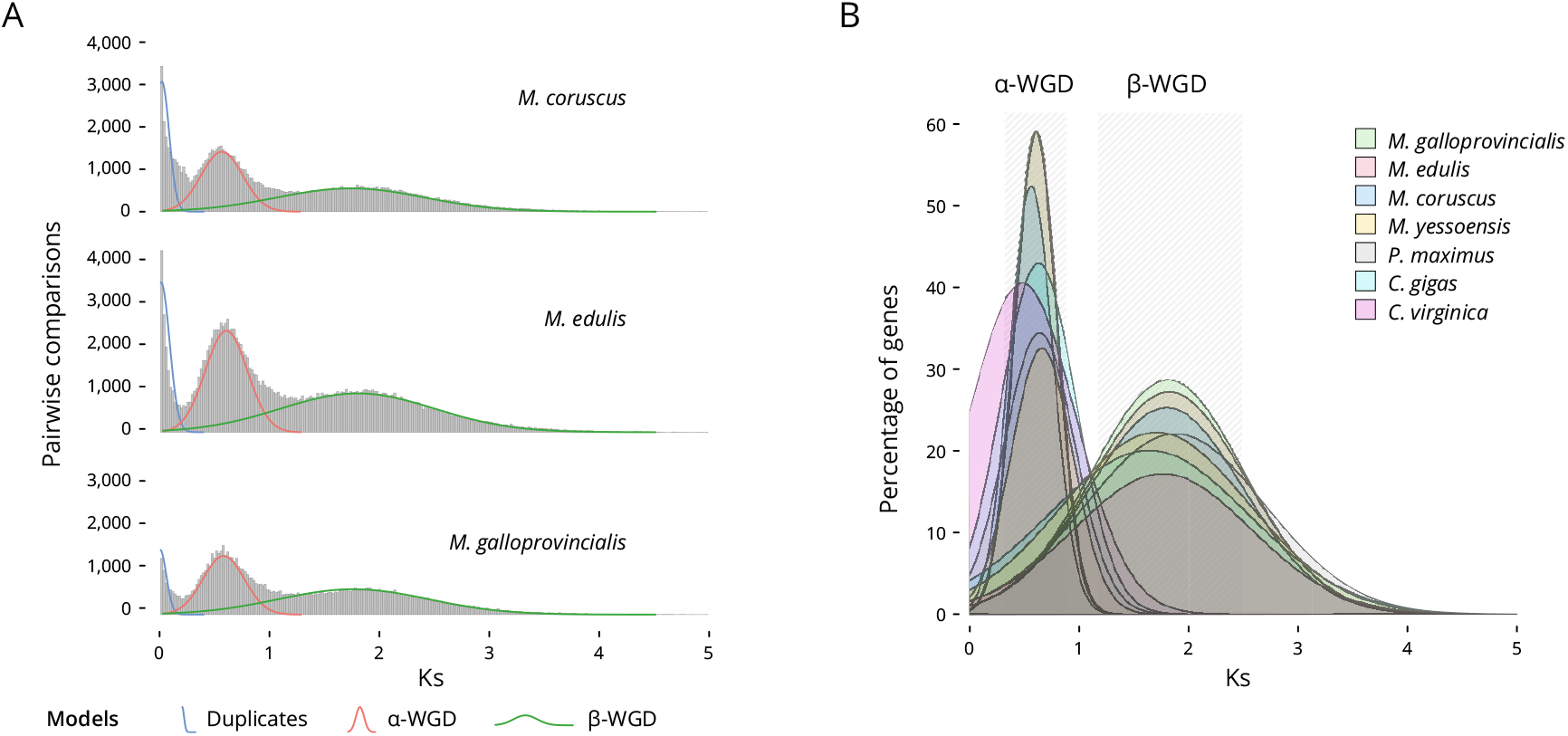
Ka and Ks analysis. **A** Distribution of the Ks values of the duplicate pairs in *M. coruscus, M. edulis* and *M. galloprovincialis*; **B** Bivalvia (class) ortholog Ks distribution and multiple WGDs. Combined Ks plot of the gene age distributions of seven species (see Tables S5 & S6). The median peaks for these plots are highlighted. Analyses of ortholog divergence indicated that these taxa diverged after their most recent WGDs.

### Identification and Functional Analysis of Positively Selected Genes

Fig. 4 shows the mean Ka/Ks ratios for each orthologs, unigene cluster, in *M. edulis* and *M. galloprovincialis* (Fig. 4A); *M. edulis* and *M. coruscus* (Fig. 4B) and *M. coruscus* and *M. galloprovincialis* (Fig. 4C). In each figure, genes clusters are colour coded to indicate groups of orthologs positively selected in both species (blue), species-specific positively selected orthologs (red), and groups of orthologs of interest (green) involved in immunity, stress response and shell formation. Collectively, the data in Fig. 4 provide a new insight into positive selection occurring in the three *Mytilus* species object of this study.

**Figure 4.**
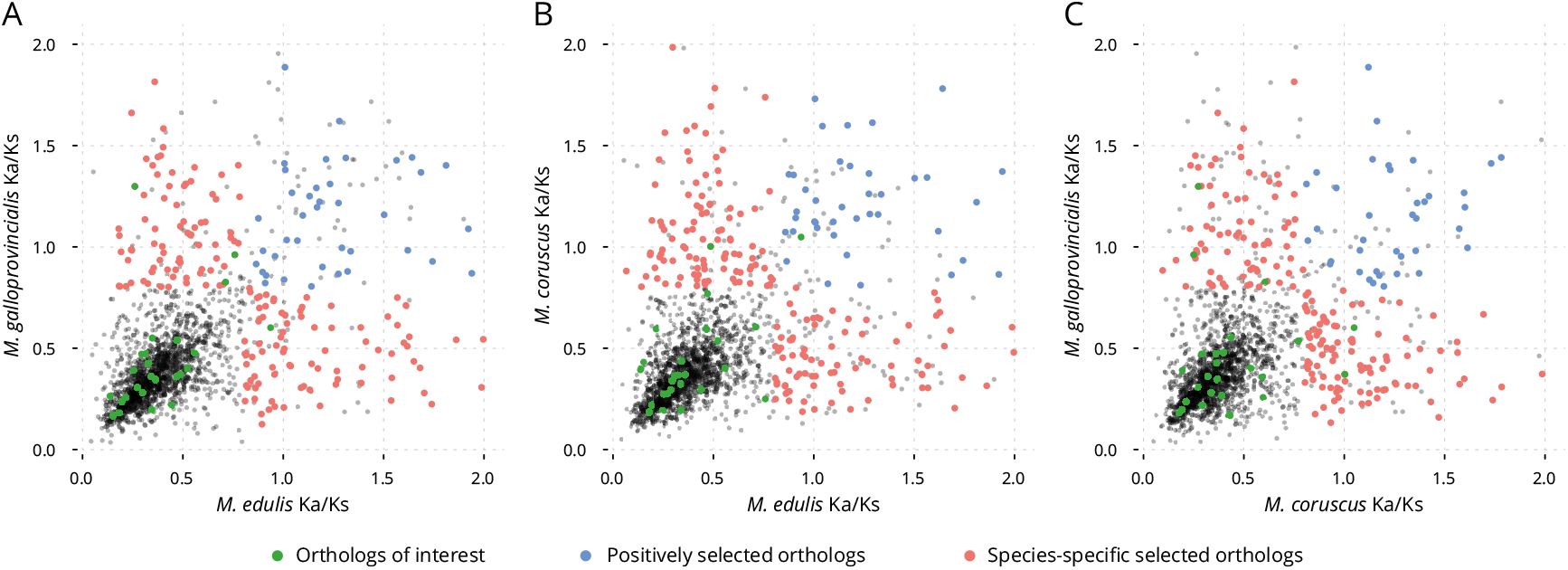
Mean Ka/Ks ratios for each orthologs cluster in the **A** *M. edulis* and *M. galloprovincialis*; **B** *M. edulis* and *M. coruscus*; **C** *M. coruscus* and *M. galloprovincialis*.

The functional analysis of positively selected orthologs has also allowed for the identification of gene duplications within assembled unigene clusters involved in key physiological processes. Here, we provide an overview of the identified functions of genes under positive selection. When *M. edulis* and *M. galloprovincialis* Ka/Ks ratios are plotted (Fig. 4A), several gene clusters can be observed to be under positive selection in both species (in blue). Of these, four were identified as contributing to immunity, stress response or shell formation: the WNT Inhibitory Factor 1 (WIF1), the nucleotide exchange factor SIL1, the kelch-like protein and the midline (MID1) protein. WIF1 contributes to several immune response functions (Capt et al., 2020), and presented a Ka/Ks value of 1.4 and 1.8 for *M. galloprovincialis* and *M. edulis* respectively. SIL1 is a protein that interacts with Hsp70 during stress responses (Fu et al., 2014), and showed a Ka/Ks value of 1.4 and 1.6 for *M. galloprovincialis* and *M. edulis*, respectively. The kelch-like protein facilitates protein binding and dimerisation (Shi et al., 2019), and presented a Ka/Ks value of 1.4 and 1.7 for *M. galloprovincialis* and *M. edulis*, respectively. Finally, the midline (MID1) protein, presenting E3 ubiquitin ligase activity (Zanchetta and Meroni, 2019), showed a Ka/Ks value of 1.4 and 1.6 for *M. galloprovincialis* and *M. edulis*, respectively.

Many of the gene clusters were also found to be under intense positive selection in *M. galloprovincialis* but under intense purifying selection in *M. edulis* or vice versa (in red). These genes are of particular interest as they indicate relatively rapid divergence between the two species. In Fig. 4, the genes with highly divergent selection are: the Glycolipid transfer protein, a ubiquitous protein characterised by their ability to accelerate the intermembrane transfer of glycolipids (Brown and Mattjus, 2007); The SGNH Hydrolase-Like Protein for which no function has been identified (Le et al., 2019); The vitellogenic carboxypeptidase-like protein, involved in key developmental processes (Sui et al., 2009). Furthermore, two unknown proteins with Ka/Ks values of 1.7 and 1.5 for *M. galloprovincialis* and Ka/Ks value of 0.2 and 0.4 for *M. edulis* have also been identified. On the other hand, gene clusters with high Ka/Ks values for *M. edulis* but low for *M. galloprovincialis* included: Mucolipin, which promotes calcium homeostasis and is involved in stress response functions (Zhang et al., 2019). The KAT8 regulatory NSL complex with developmental and cellular homeostasis function (Radzisheuskaya et al., 2021). The TolB-like protein, involved in a tol-dependent translocation system (Carr et al., 2000). The RING finger protein 170, which mediates the ubiquitination and degradation of inositol 1,4,5-trisphosphate receptors, and it is involved in immune response functions (Song et al., 2019) and finally, Fibropellin, a cell adhesion protein (Nie et al., 2020). Only a limited number of gene clusters appear to be under positive selection amongst those commonly used in immunity, stress response and shell formation comparative studies (in green), and the majority of the genes clusters showed negative selection with the exception of four, which resulted to be all involved in immune response pathways (Gerdol and Venier, 2015). Of these, three were positively selected in *M. galloprovincialis* and conserved in *M. edulis* (membrane-bound C-type lectin, Galectin 3, MAP kinase 4-like) and one was positively selected in *M. edulis* and conserved in *M. galloprovincialis* (TNF ligand-like 2).

The comparison of Ka/Ks values between *M. coruscus* and *M. edulis* (Fig. 4B) shows a similar picture to that of *M. edulis* and *M. galloprovincialis* (Fig. 4A). Positively selected orthologs in both species (with Ka/Ks values ranging from 1.3 to 1.9) include: the nucleotide exchange factor SIL1; the archease-like protein, related to stress response functions (Auxilien et al., 2007); the midline (MID1) protein and the thiosulfate/3-mercaptopyruvate sulfotransferase protein involved in developmental and stress response functions (Mao et al., 2011). The ortholog clusters positively selected in *M. coruscus* but conserved in *M. edulis* (Ka/Ks values from 1.6 to 2.0 and from 0.3 to 0.7 respectively) include: the purine-nucleoside phosphorylase, which encodes an enzyme which reversibly catalyses the phosphorolysis of purine nucleosides (Stoeckler et al., 1997); the palmitoyl-protein thioesterase, which facilitates the morphological development of neurons and synaptic function in mature cells (Koster, 2019), and the ADAR protein which is an RNA-binding protein and has antiviral immunity in marine molluscs (Green et al., 2015). Two further proteins with unknown associated functions and with Ka/Ks values of 1.8 and 1.7 for *M. coruscus* and of 0.3 and 0.5 for *M. edulis* were also identified. Similarly, orthologs positively selected in *M. edulis* but conserved in *M. coruscus* were functionally characterised and resolved to be the same as those described for *M. edulis* and *M. galloprovincialis* (Fig. 4A). Only two gene clusters showed positive selection amongst those of physiological interest (Fig. 4B, in green). C-type lectin 7 (immunity) was positively selected in *M. coruscus* with a Ka/Ks value of 1.0 and 0.4 for *M. coruscus* and *M. edulis*, respectively; while TNF ligand-like 2 (immunity) is presenting positive selection in both species, with Ka/Ks values of 1.0 and 0.9 for *M. coruscus* and *M. edulis*, respectively.

In the *M. galloprovincialis* and *M. coruscus* comparison (Fig. 4C), positively selected proteins in both species (in blue) include: the nucleotide exchange factor SIL1, the importin-7 protein, involved in nuclear import of histones and the homeobox protein cut-like (CUTL), involved in cell-cell adhesion interactions that are required for normal development (Pérez-Parallé et al., 2005). The ortholog clusters positively selected in *M. galloprovincialis* (in red) but conserved in *M. coruscus* (Ka/Ks values from 1.4 to 1.8 and from 0.2 to 0.7 respectively) are the same as described in *M. edulis* and *M. galloprovincialis* (Fig. 4A). The ortholog clusters positively selected in *M. coruscus* (in red) but conserved in *M. galloprovincialis* (Ka/Ks values from 1.6 to 1.9 and 0.3 to 0.4 respectively) are again the same as described in *M. coruscus* and *M. edulis* (Fig. 4B). The proteins positively selected for *M. galloprovincialis* and conserved for *M. coruscus* (Fig. 4C, in green) are: the membrane-bound C-type lectin (immunity) with a Ka/Ks value of 1.3 and 0.3 for *M. galloprovincialis* and *M. coruscus*, respectively. The galectin 3 protein (immunity) with a Ka/Ks value of 1.0 and 0.3 for *M. galloprovincialis* and *M. coruscus*, respectively. And the MAP kinase 4-like protein (immunity) with a Ka/Ks value of 0.8 and 0.6 for *M. galloprovincialis* and *M. coruscus*, respectively. Finally, two genes showed positive selection among those of physiological interest (in green) in favour of *M. coruscus*, and conserved for *M. galloprovincialis*. The TNF ligand-like 2 (immunity) with a Ka/Ks value of 1.0 and 0.6 for *M. coruscus* and *M. galloprovincialis*, respectively. And C-type lectin 7 (immunity) with a Ka/Ks value of 1.0 and 0.4 for *M. coruscus* and *M. galloprovincialis*, respectively.

The vast majority of our orthologs of interest (in green) selected from the literature have not shown a substantial number of proteins under positive selection for genes related to immunity, stress response, and shell formation.

For completeness, all genes involved in immunity, stress response and shell formation under positive selection in any of the three species examined here, were identified and grouped by species (Table S7). For *M. galloprovincialis*, 6, 10 and 3 genes related to immunity, stress response and shell formation, respectively were detected. For *M. edulis*, 13, 6 and 4 genes related to immunity, stress response and shell formation, respectively were detected, and for *M. coruscus*, 10, 5 and 2 genes related to immunity, stress response and shell formation, respectively were detected.

## DISCUSSION

### Reference Genome and Whole Genome Duplication

Mussels are also known as poor man’s shellfish as they are inexpensive and abundant. These features have perhaps contributed to a relative neglect in the investigation of this species’ genomic structural variation, and whether such structural changes can play a significant role in their evolution and ecological adaptations (Davison and Neiman, 2021). In the wild, mussels thrive on rocks and stones along the coast, but the majority of mussels consumed are farmed in coastal waters providing food security and employment opportunities to a multitude of fragile coastal communities worldwide (Smaal et al., 2019). Similarly to several other molluscs classes, genomic research into *M. edulis* has been hampered by the lack of a reference genome. This bottleneck is historically linked with the technical difficulties in extracting high molecular weight genomic DNA from Molluscan tissues and thus, allow for long reads sequencing techniques to be successfully applied (Davison and Neiman, 2021). In addition, the relatively large genome size and a high level of heterozygosity further complicates the assembly of high-quality reference genomes in the phylum.

With the aim of shedding new light onto the genomic structure and evolution of the class Bivalvia, we sequenced the blue mussel genome and we introduced the first evidence of WGD events in the Mytilidae family and in Bivalvia more generally. Finally, we identify genes within key physiological pathways under significant positive selection. Taken together, our results provide new insights into the Mytilidae family genome structure and introduce new genomic resources for the investigation of Bivalves evolution, population genetics and for future selective breeding activities. The genome was assembled into 3,339 scaffolds with a total length of 1.83 Gb, a GC content of 32.17% and a scaffold N_50_ of 1.10 Mb. In addition, we found 1.03 Gb (56.33% of the assembly) of repeat content, 69,246 protein-coding genes, 132 rRNAs and a heterozygosity of 0.48% (Table 1). The results are equivalent with the other *Mytilus* genomes: Genome size between 1.90 Gb and 1.28 Gb, and repetitive sequences between 52.83% and 58.56% (Gerdol et al., 2020; Li et al., 2020; Murgarella et al., 2016; Yang et al., 2021). In addition, transcriptomic data and the derived gene models are comparable with the other available Mytilidae transcriptomes. Phylogeny of the *Mytilus* (based on the mitochondrial genomes) confirms the position of *M. edulis* in the genus, with the *M. edulis* and *M. galloprovincialis* (sympatric species) separate from *M. trossulus* and *M. coruscus* (which group with *M. californianus*; Fig. 2B).

Our analysis provides, for the first time, genomic evidence for paleopolyploidy in the class Bivalvia. Combining our gene age distribution and phylogenomic analyses, we found evidence for two significant, episodic bursts of gene duplication. While some of these duplication events may be caused by other processes of gene duplication, they are compatible with WGDs observed using similar methods in plants (Badouin et al., 2017) and animals (Berthelot et al., 2014; Li et al., 2018). Ks analysis showed that an ancient WGD event and a more modern WGD event occurred before the divergence of the Bivalvia. This explains why bivalves, and molluscs more generally, present large genomes (Davison and Neiman, 2021). The genomic information for *M. edulis* presented here, will help clarify the evolutionary processes in Bivalvia species and contribute to improving the understanding of the physiological and morphological diversity of Bivalvia species. Our discovery of WGDs in the ancestry of Bivalvia raises questions about the role of gene and genome duplication in their evolution. After duplication, the most likely fate of duplicated genes is the loss of one of the duplicates through non-functionalisation that occurs by accumulation of deleterious mutations (Nei and Roychoudhury, 1973; Takahata and Maruyama, 1979). While common after WGD, gene loss could however play a key role in speciation (Lynch, 2000), through a process known as divergent resolution (Taylor et al., 2001). In addition, duplicated genes may also be retained in two copies (Ohno, 1970) and either specialise by the partitioning of ancestral gene functions (*i*.*e*. sub-functionalisation) or by the acquisition of a novel function (*i*.*e*. neo-functionalisation).

Incomplete genetic data (draft genomes and transcriptomes), as well as reduced datasets (Enzymes, RAD, or EST), made it impossible to correctly detect WGDs and duplicated genes in Bivalvia, before now. In the absence of complete genomes and the full picture of WGD events, duplicated sequences are often overlooked or wrongly interpreted. This can lead to artefacts such as high heterozygosity (Vendrami et al., 2020), pseudogenes, and a rapid rate of gene acquisition and loss (Gerdol et al., 2020). The discovery of several events of WGD in the Bivalvia phylogeny suggests the prospect that large-scale duplications are consistent with the evolution of novelty and diversity in the physiology of mass spawners like Bivalvia. However, dating of such events remains difficult due to the lack of annotated genomes deeper in the phylogeny, which still is a priority to fully elucidate molluscan evolution.

### Identification and Functional Analysis of Positively Selected Genes

The functional analysis of positively selected orthologs has allowed us to compare our results with studies related to the identification of gene involved in key physiological processes. When identifying gene clusters under positive selection in both species (blue dots in Fig. 4; *M. galloprovincialis*-*M. edulis, M. coruscus*-*M. edulis*, and *M. galloprovincialis*-*M. coruscus*), we find the predominant functions for those genes are mainly related to immunity, stress responses and developmental processes. Our results agree with past studies confirming that genes related to immunity are under selection in multiple lineages, likely via adaptive evolution mechanisms linked to host-pathogens co-evolution (Oliver et al., 2010). The stress response genes presenting positive selection are related or are interacting with Hsp proteins (SIL1 and the archease-like protein). Since the marine environment has considerable concentration of bacteria and viruses, molluscs depend on cellular and molecular mediated immune responses that help them to survive under challenging conditions (Pourmozaffar et al., 2020). That is why filter-feeding animals such as bivalves rely on the intervention of shock proteins which synthesis depends on environmental stressful conditions such as temperature, salinity, hypoxia, heavy metal, and infectious pathogens (Wan et al., 2012). Since we are comparing three species adapted to different environmental conditions, our results suggest that stress genes are evolving independently to adapt to specific environmental conditions.

Genes presenting intense positive selection in one species but intense purifying selection in the others are of interest because they indicate rapid divergence between species. Once again, the three species have their maximum Ka/Ks values in genes related to developmental processes, immunity, and stress response. Overall, genes identified as being under positive selection in this study, are consistent with the defence system of bivalves depending on the innate immune response against stressful conditions such as environmental stressors, pollution and disease outbreaks.

The identification of all the genes involved in immunity, stress response and shell formation under positive selection in any of the three species (Table S7) has provided us with relevant information that could be used in future studies to identify markers for future comparative physiology and evolution studies. Our results for *M. galloprovincialis* have shown a considerable amount of stress response proteins (10 proteins) under positive selection compared to the other two species. A significant amount of those stress response genes have documented roles in heat tolerance or direct associations to heat-stress responses, *e*.*g*. zinc finger MYM-type protein 2-like (ZMYM2), mitogen-activated protein kinase 6 (MAPK6), heat shock protein 22 (HSPB8). This is also supported by past studies (Saarman et al., 2017; Popovic and Riginos, 2020) were genomic functions previously linked to divergent temperature adaptation reflected accelerated molecular divergence between warm-adapted *M. galloprovincialis* and cold-adapted congeners, such as *M. edulis*. Molecular divergence of *M. galloprovincialis* is consistent with warm-temperature adaptation demonstrated by physiological studies. *M. galloprovincialis* also has more positive selection in stress response proteins related to heavy metal detection, transport and metal binding (*e*.*g*. arylesterase / paraoxonase, pyruvate dehydrogenase E1 component alpha subunit, inositol polyphosphate 1-phosphatase, Solute carrier family 12), than *M. edulis* and *M. coruscus. M. edulis* and *M. galloprovincialis* presented positively selected shell formation proteins, in the EF-hand domains, which appears to be evolving faster in the two species, albeit in different gene clusters: EF-hand domain protein, EF-Hand, calcium-binding site for *M. edulis* and EF-hand calcium-binding domain-containing protein for *M. galloprovincialis*. In bivalves, the Ca^++^ binding EF hand domains include a Calmodulin-like protein (CaLP), a multifunctional calcium sensor that belongs to a new member of the CaM (cell adhesion molecules) superfamily, localised in the organic layer sandwiched between nacre and prismatic (aragonite) layer (calcite) in *Pinctada fucata* (Yan et al., 2007). Studies have shown that CalP might be involved in the growth of nacre layer and prismatic layer (Feng et al., 2017). Our results suggest and support past studies indicating that closely related bivalves use different secretory repertoires to construct their shell (Peterson et al., 2008) which might lead to positive selection at a gene level as reflected in our results. Also, shell dissolution and decreased shell growth caused by ocean acidification have been described in marine bivalves (Melzner et al., 2011) forcing the need for a fast environmental adaptation. Taking in account current alterations in precipitation patterns as well as stronger and more frequent heat waves and fluctuating sea surface salinities (Steeves et al., 2018), our results suggest that *M. galloprovincialis* appears to be better equipped than *M. edulis* and *M. coruscus* to adapt to higher temperatures, aquatic toxicity, and contamination.

The recruitment, settlement, and grow-out phase of bivalve aquaculture and more precisely, in *Mytilus* spp. is strongly dependant with the environmental conditions. Therefore, the implications of climate change are not restricted to wild populations. Strong changes in local environmental conditions may limit production and force the relocation of grow-out sites to suitable areas. Thinner and weaker shells will facilitate their rupture during transportation and increase losses due to predation. Genomic selection studies and identification of molecular markers can favour the development of genetically improved lines for multiple traits and facilitate the management of genetic variability. The development of high-quality assembled genomes, as provided by the current research, will favour the identification of genomic regions linked to traits responsible for environmental resilience, which will support the long-term sustainable management and exploitation of the species.

## DATA AVAILABILITY STATEMENT

The raw sequencing reads of all libraries are available from EBI/ENA via the accession numbers ERR4296957-ERR4296958 (long reads), ERR4296954-ERR4296955 (short reads) and ERR4172341 (RNA-seq). The assembled genome (MEDL1; GCA 905397895.1) is available in EBI with the accession numbers ERS4576331, project PRJEB38403.

## ETHICS STATEMENT

This work was approved by the University of Stirling Ethics Committee (Animal Welfare and Ethics Review Board). Animal handling and collection in this study was carried out following its approved guidelines and regulations.

## AUTHOR CONTRIBUTIONS

S.C. sourced the funding, co-developed the conceptual idea, supervised the findings of this work and edited the manuscript. M.B. co-developed the concept with S.C., curated all the computational and bioinformatic elements of the study and edited the manuscript. A.C.F. sourced and prepared the biological material, wrote the first draft of the manuscript and conducted the functional analysis of all orthologs, A.D. extracted the RNA and edited the final manuscript.

