## Supplementary Figure S1 and Supplementary Tables S1-S7 for "Evidence of multiple genome duplication events in *Mytilus* evolution"

### ABSTRACT

Molluscs remain one significantly under-represented taxa amongst available genomic resources, despite being the second-largest animal phylum and the recent advances in genomes sequencing technologies and genome assembly techniques. With the present work, we want to contribute to the growing efforts by filling this gap, presenting a new high-quality reference genome for *Mytilus edulis* and investigating the evolutionary history within the Mytilidae family, in relation to other species in the class Bivalvia.
Here we present, for the first time, the discovery of multiple whole genome duplication events in the Mytilidae family and, more generally, in the class Bivalvia. In addition, the calculation of evolution rates for three species of the Mytilinae subfamily sheds new light onto the taxa evolution and highlights key orthologs of interest for the study of *Mytilus* species divergences.
The reference genome presented here will enable the correct identification of molecular markers for evolutionary, population genetics, and conservation studies. Mytilidae have the capability to become a model shellfish for climate change adaptation using genome-enabled systems biology and multi-disciplinary studies of interactions between abiotic stressors, pathogen attacks, and aquaculture practises.

### SUPPLEMENTARY FIGURE

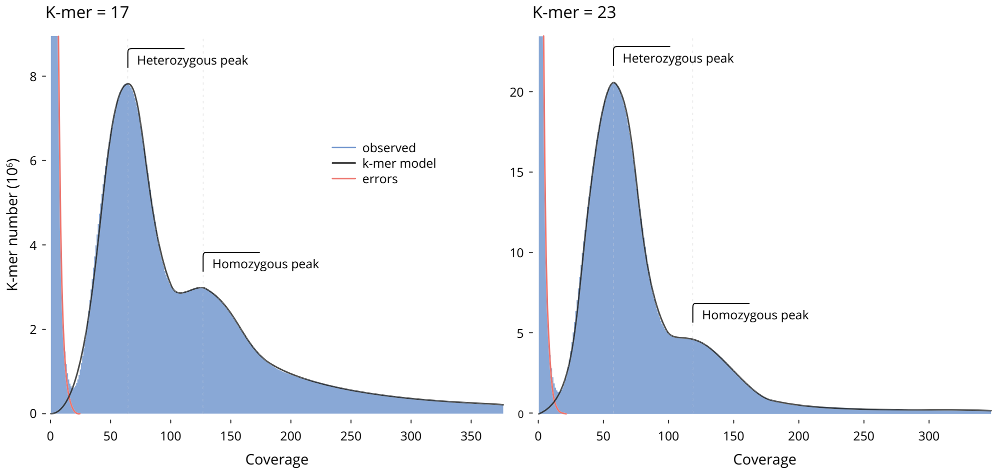


**Figure 1.** The k-mer distribution used for the estimation of genome size. The heterozygous and homozygous peaks of k-mer depth are clearly markers, suggesting a high-complexity genome. **A** The 17-mer distribution. Predicted genome size, 1,010,184,781 nt; **B** The 23-mer distribution. Predicted genome size, 1,096,306,163 nt.

### SUPPLEMENTARY TABLES

**Table 1.** Sequencing data, summary statistics. ∗ Estimation based on *M. galloprovincialis* and *M. coruscus* genomes size.

| Category | Number/length |
|---|---|
| Total number of long reads | 15,945,130 |
| Total number of bases | 111,654,433,463 |
| Mean length | 7,002 nt |
| Maximum read length | 672,255 nt |
| Coverage* | 64x |
| Total number of PE short reads | 652,465,784 |
| Total number of bases | 195,739,735,200 |
| Read length | 150 nt |
| Coverage* | 113x |
| Total number of RNA-seq PE short reads | 50,593,080 |
| Total number of bases | 15,177,924,000 |
| Read length | 150 nt |
| Coverage* | 9x |

**Table 2.** RepeatMasker statistics. ∗ repeats fragmented by insertions or deletions have been counted as one element. † LTR_Finder results: 255,413 LTR pairs over 4,932 regions and 151,703,353 bp.

| Element | Number of elements* | Length occupied | Percentage of sequence |
|---|---|---|---|
| SINEs | 10,261 | 2,030,467 bp | 0.11% |
| ALUs | 0 | 0 bp | 0.00% |
| MIRs | 0 | 0 bp | 0.00% |
| LINEs | 343,643 | 130,980,160 bp | 7.17% |
| LINE1 | 11,902 | 5,537,561 bp | 0.30% |
| LINE2 | 7,970 | 3,762,443 bp | 0.21% |
| L3/CR1 | 6,535 | 4,167,698 bp | 0.23% |
| LTR elements† | 4,932 | 151,703,353 bp | 8.30% |
| DNA elements | 99,267 | 25,504,172 bp | 1.40% |
| hAT-Charlie | 2,603 | 433,449 bp | 0.02% |
| TcMar-Tigger | 0 | 0 bp | 0.00% |
| Unclassified | 2,976,814 | 707,039,397 bp | 38.70% |
| Small RNA | 133 | 25,717 bp | 0.00% |
| Satellites | 2,252 | 400,532 bp | 0.02% |
| Simple repeats | 224,100 | 10,280,290 bp | 0.56% |
| Low complexity | 50,703 | 2,473,133 bp | 0.14% |
| Total repeats | | 1,029,206,554 bp | 56.33% |

**Table 3.** Summary of annotation results for *M. edulis* gene models using a range of databases. $*$ InterPro covers 12 databases (CDD-3.17,Coils-2.2.1, Gene3D-4.2.0, Hamap-2020_01, MobiDBLite-2.0, PANTHER-14.1, PRINTS-42.0, ProSitePatterns-2019_11, ProSiteProfiles-2019_11, SFLD-4, SMART-7.1, SUPERFAMILY-1.75, TIGRFAM-15.0).

| Database | Number annotated |
|---|---|
| PfamA | 61,453 |
| InterPro* | 48,772 |
| SwissProt | 11,211 |
| KEGG | 51,091 |
| GO | 31,620 |
| All | 9,089 |
| Total | 69,246 |

**Table 4.** Mytilinae (subfamily) mitochondrial genomes.

| Accession | Genome size | Species name | Common name |
|---|---|---|---|
| NC_018362.1 | 16,014 bp | *Perna viridis* | Asian green mussel |
| NC_044131.1 | 16,253 bp | *Septifer bilocularis* | - |
| NC_044128.1 | 17,582 bp | *Crenomytilus grayanus* | - |
| NC_030633.1 | 16,765 bp | *Mytilus chilensis* | Chilean mussel |
| NC_028706.1 | 18,145 bp | *Limnoperna fortunei* | Golden mussel |
| NC_026288.1 | 18,415 bp | *Perna perna* | Brown mussel |
| NC_024733.1 | 16,642 bp | *Mytilus coruscus* | Hard-shell mussel |
| NC_006886.2 | 16,744 bp | *Mytilus galloprovincialis* | Mediterranean mussel |
| NC_006161.1 | 16,740 bp | *Mytilus edulis* | Blue mussel |
| NC_007687.1 | 18,652 bp | *Mytilus trossulus* | Bay mussel |
| NC_015993.1 | 16,730 bp | *Mytilus californianus* | California mussel |

**Table 5.** Bivalvia (class) genome where Ka & Ks estimations were possible: All exhibit evidences of $\alpha$-WGD and $\beta$-WGD. $*$ this study.

| Species | Peak/Mean ($\alpha$-WGD) | Std. dev. ($\alpha$-WGD) | Peak/Mean ($\beta$-WGD) | Std. dev. ($\beta$-WGD) |
|---|---|---|---|---|
| *M. edulis** | 0.6132 | 0.1939 | 1.8196 | 0.7103 |
| *M. galloprovincialis* | 0.6010 | 0.1934 | 1.8109 | 0.7011 |
| *M. coruscus* | 0.5635 | 0.1996 | 1.7977 | 0.7082 |
| *C. gigas* | 0.6299 | 0.3699 | 1.6350 | 0.4601 |
| *C. virginica* | 0.4787 | 0.4921 | 1.7627 | 0.3479 |
| *M. yessoensis* | 0.6612 | 0.2278 | 1.7064 | 0.4722 |
| *P. maximus* | 0.6392 | 0.2759 | 1.8794 | 0.4541 |

**Table 6.** Bivalvia (class) genome and availability of gene models and annotations. ∗ this study.

| Assembly | Reference | Species name | Common name | Gene Models |
|---|---|---|---|---|
| GCA_014843695.1 | ASM1484369v1 | *Archivesica marissinica* | Deep-sea clam | - |
| GCA_004382765.1 | QAU_Acon_1.1 | *Argopecten irradians* concentricus | Bay scallop | - |
| GCA_004382745.1 | QAU_Airr_1.1 | *Argopecten irradians* irradians | Bay scallop | - |
| GCA_002080005.1 | Bpl_v1.0 | *Bathymodiolus platifrons* | Cold seep mussel | - |
| GCA_001632725.1 | ASM163272v1 | *Corbicula fluminea* | Asian clam | - |
| GCF_902806645.1 | cgigas_uk_roslin_v1 | *Crassostrea gigas* | Pacific oyster | yes |
| GCA_005518195.2 | NWPU_Cgig_v2 | *Crassostrea gigas* | Pacific oyster | - |
| GCF_000297895.1 | oyster_v9 | *Crassostrea gigas* | Pacific oyster | - |
| GCA_000297895.2 | ASM29789v2 | *Crassostrea gigas* | Pacific oyster | - |
| GCA_011032805.1 | ASM1103280v1 | *Crassostrea gigas* | Pacific oyster | - |
| GCA_015776775.1 | ASM1577677v1 | *Crassostrea hongkongensis* | Hong Kong oyster | - |
| GCA_002022765.1 | C_virginica_1.0 | *Crassostrea virginica* | Eastern oyster | - |
| GCA_002022765.3 | C_virginica-2.0 | *Crassostrea virginica* | Eastern oyster | - |
| GCF_002022765.2 | C_virginica-3.0 | *Crassostrea virginic*a | Eastern oyster | yes |
| GCA_012932295.1 | ASM1293229v1 | *Cyclina sinensis* | Venus clam | - |
| GCA_000806325.1 | ASM80632v1 | *Dreissena polymorph*a | Zebra mussel | - |
| GCA_007657795.1 | UV_Dro_v1.1 | *Dreissena rostriformis* | Quagga mussel | - |
| GCA_003130415.1 | ASM313041v1 | *Limnoperna fortunei* | Golden mussel | - |
| GCA_008271625.1 | LuRhyn_1.0 | *Lutraria rhynchaena* | Snout Otter Clam | - |
| GCA_016163765.1 | CUHK_oyster_2.0 | *Magallana hongkongensi*s | Hong Kong oyster | - |
| GCA_015947965.1 | ASM1594796v1 | *Margaritifera margaritifera* | Freshwater pearlshell mussel | - |
| GCA_016617855.1 | ASM1661785v1 | *Megalonaias nervosa* | Washboard | - |
| GCA_014805675.1 | ASM1480567v1 | *Mercenaria mercenaria* | Northern quahog | - |
| GCF_002113885.1 | ASM211388v2 | *Mizuhopecten yessoensis* | Yesso scallop | yes |
| GCA_002113885.1 | ASM211388v1 | *Mizuhopecten yessoensis* | Yesso scallop | - |
| GCA_002080025.1 | Mph_v1.0 | *Modiolus philippinarum* | Philippine horse mussel | - |
| GCA_017311375.1 | Mcoruscus_HiC | *Mytilus coruscu*s | Hard-shell mussel | - |
| GCA_011752425.2 | MCOR1.1 | *Mytilus coruscus* | Hard-shell mussel | yes |
| GCA_011752425.1 | MCOR1 | *Mytilus coruscus* | Hard-shell mussel | - |
| GCA_905397895.1* | MEDL1 | *Mytilus edulis* | Blue mussel | yes |
| GCA_900618805.1 | MGAL_10 | *Mytilus galloprovincialis* | Mediterranean mussel | yes |
| GCA_000715055.1 | mussel1.0 | *Mytilus galloprovincialis* | Mediterranean mussel | - |
| GCA_001676915.1 | ASM167691v1 | *Mytilus galloprovinciali*s | Mediterranean mussel | yes |
| GCA_903981925.1 | v081 | *Ostrea lurida* | Olympia oyster | - |
| GCA_902825435.1 | PGEN-v1.0 | *Panopea generosa* | Pacific geoduck | - |
| GCF_902652985.1 | xPecMax1.1 | *Pecten maximus* | King scallop | yes |
| GCA_002216045.1 | PinMar1.0 | *Pinctada imbricata* | Akoya pearl oyster | - |
| GCA_016161895.1 | ASM1616189v1 | *Pinna nobilis* | Noble penshell | - |
| GCA_016746295.1 | UT_Pstr_1.0 | *Potamilus streckersoni* | Brazos heelsplitter | - |
| GCA_009026015.1 | ASM902601v1 | *Ruditapes philippinarum* | Manila clam | - |
| GCA_003671525.1 | Sgl1.0 | *Saccostrea glomerata* | Sydney rock oyster | - |
| GCA_007844125.1 | ASM784412v1 | *Sinonovacula constrict*a | Chinese razor clam | - |
| GCA_009762815.1 | ASM976281v1 | *Sinonovacula constricta* | Chinese razor clam | - |
| GCA_013375625.1 | ASM1337562v1 | *Tegillarca granosa* | Blood clam | - |
| GCA_003401595.1 | ASM340159v1 | *Venustaconcha ellipsiformis* | Ellipse | - |

**Table 7.** Genes involved in immunity, stress response and shell formation under positive selection in *M. galloprovincialis*, *M. edulis* and *M. coruscus*.

| Species | Category | ClusterID | Gene Names / Function (Ortholog cluster) | Ka/Ks | Ortholog reference |
|---|---|---|---|---|---|
| *M. galloprovincialis* | Immunity | 9813 | NIT1<br>Potential tumour suppressor in tumour progression | 2.5 | Lin et al. (2018) |
| *M. galloprovincialis* | Stress | 6307 | Arylesterase / Paraoxonase<br>Confer resistance to organophosphate toxicity | 1.4 | Bonacci et al. (2004) |
| *M. galloprovincialis* | Stress | 1976 | Serine/threonine kinase 17<br>Cellular processes, proliferation, apoptosis, and differentiation. Abiotic stress | 1.4 | Zorina et al. (2014) |
| *M. galloprovincialis* | Stress | 8246 | Peptidase inhibitor 16<br>Cardiac stress response | 1.4 | Regn et al. (2016) |
| *M. galloprovincialis* | Shell formation | 18160 | EF-hand Ca2+-binding domain 6<br>CaLP has two Ca2+-binding EF hand domains.Growth of nacre-prismatic layer | 1.2 | Feng et al. (2017) |
| *M. galloprovincialis* | Immunity | 19933 | Signal transducer-activator of transcription 5B<br>Mediate the signaling of cytokines and a number of growth factors | 1.1 | Yu et al. (2019) |
| *M. galloprovincialis* | Stress | 1594 | Pyruvate dehydrogenase E1 alpha subunit<br>Reduces OXPHOS and oxygen consumption | 1.1 | Zimmer et al. (2016) |
| *M. galloprovincialis* | Stress | 973 | DNA mismatch repair protein MSH6<br>Responsive to oxidative stress and protection against ROS and DNA damage | 1.1 | Pinheiro et al. (2021) |
| *M. galloprovincialis* | Stress | 25447 | zinc finger MYM-type protein 2-like<br>Significant correlations with salinity, temperature, As, Cd or lindane | 1.1 | Baillon et al. (2015) |
| *M. galloprovincialis* | Shell formation | 12897 | Ca2+ transporting ATPase, plasma membrane<br>Catalyse the hydrolysis of ATP coupled with the transport of calcium | 1 | Sillanpää et al. (2018) |
| *M. galloprovincialis* | Stress | 14603 | Inositol polyphosphate 1-phosphatase<br>Mg2+ dependent of inositol monophosphatase-like domain | 1 | Bialojan and Takai (1988) |
| *M. galloprovincialis* | Immunity | 14873 | Galectin-4<br>Regulators of immune cell homeostasis | 1 | Vasta et al. (2015) |
| *M. galloprovincialis* | Immunity | 3134 | PRRT1<br>Proline-rich transmembrane protein 1 | 1 | Marin et al. (2000) |
| *M. galloprovincialis* | Shell formation | 14056 | Metalloproteinase inhibitor 3<br>Ligament-specific protein | 0.9 | Kubota et al. (2017) |
| *M. galloprovincialis* | Stress | 1893 | Solute carrier family 12<br>Transport endogenous-exogenous substances. Potassium/chloride transporters | 0.9 | Xun et al. (2020) |
| *M. galloprovincialis* | Immunity | 7356 | Leucine-rich repeat domain superfamily<br>Development, growth, and responses to abiotic and biotic stresses | 0.9 | Wang et al. (2017) |
| *M. galloprovincialis* | Stress | 3258 | Heat shock protein 22 HSPB8<br>Protecting cells, folding of nascent peptides, and responding to stress | 0.9 | Zhang et al. (2010) |
| *M. galloprovincialis* | Stress | 5345 | Mitogen-activated protein kinase 6 MAPK<br>involved in the regulation of Hsp expression in blue mussels | 0.9 | Anestis et al. (2007) |
| *M. galloprovincialis* | Immunity | 5490 | LRP1B<br>Low-densitylipoproteinreceptor-related protein | 0.9 | Liu et al. (2014b) |
| *M. edulis* | Immunity | 1359 | E3 ubiquitin-protein ligase mind-bomb (MIB2)<br>Antiviral immunity | 2.4 | Chen et al. (2015) |
| *M. edulis* | Stress | 5390 | Mucolipin, Polycystin cation channel<br>Calcium homeostasis | 2 | Jiao et al. (2019) |
| *M. edulis* | Immunity | 13969 | RNF170, RING finger protein 170<br>Ubiquitination and degradation of inositol 1,4,5-trisphosphate receptors | 1.7 | Song et al. (2019) |
| *M. edulis* | Stress | 9109 | Fibropellin-1<br>Hypoxia responsive gene | 1.7 | Nie et al. (2020) |
| *M. edulis* | Stress | 10101 | Carbohydrate-binding WSC<br>Plasma membrane sensor for surface stress | 1.5 | Oide et al. (2019) |
| *M. edulis* | Immunity | 4324 | Lactamase_B<br>Drug resistance among gram-negative bacteria | 1.3 | Singh et al. (2019) |
| *M. edulis* | Stress | 4307 | polycystin<br>Calcium homeostasis | 1.3 | Wang et al. (2020) |
| *M. edulis* | Immunity/stress | 9741 | Papain-like cysteine peptidase superfamily<br>Prevent unwanted protein degradation | 1.2 | Liu et al. (2018) |
| *M. edulis* | Immunity | 7225 | Peptidase M12B<br>Cell adhesion, signaling, cell-cell fusion, and cell-cell interactions | 1.2 | Rubin et al. (2014) |
| *M. edulis* | Stress | 7276 | ATP-dependent metalloprotease (YME1L)<br>Stress-sensitive mitochondrial protease | 1.2 | Rainbolt et al. (2015) |
| *M. edulis* | Immunity | 14877 | P-loop - nucleoside triphosphate hydrolase<br>This domain shows a high specificity for pathogens and parasites | 1.1 | Arivalagan et al. (2017) |
| *M. edulis* | shell formation | 15346 | EF-hand Ca2+-binding domain<br>CaLP has two Ca2+-binding EF hand domains(Growth of nacre-prismatic layer) | 1.1 | Feng et al. (2017) |
| *M. edulis* | Immunity | 2105 | C-type lectin superfamily 17 member A<br>Mediate crucial cellular functions during immunity and homeostasis | 1 | Kerscher et al. (2013) |
| *M. edulis* | Immunity | 10990 | Fibrinogen-like protein A<br>Immunepattern-recognition receptors. | 1 | Gorbushin and Iakovleva (2011) |
| *M. edulis* | Immunity | 20412 | FYVE, RhoGEF and PH domain<br>Signal transduction | 0.9 | Perrier et al. (2020) |
| *M. edulis* | shell formation | 5702 | Perlucin-like protein<br>Ca2+-dependent carbohydrate binding activity | 0.9 | Blank et al. (2003) |
| *M. edulis* | Immunity | 22962 | C1q-related factor<br>Pattern recognition receptors. Activates innate immune response | 0.9 | Jiang et al. (2020) |
| *M. edulis* | Immunity | 11253 | nicotinic acetylcholine receptor alpha-7<br>Regulates immune response through the neuroendocrine-immune system | 0.9 | Jiao et al. (2019) |
| *M. edulis* | Immunity | 19695 | TRIM56, tripartite motif-containing protein 56<br>virus-inducible E3 ubiquitin ligase that restricts pestivirus infection | 0.9 | Liu et al. (2014a) |
| *M. edulis* | Immunity | 11784 | HMCN, hemicentin<br>Immune recognition, signaling and regulation.insulin peptide receptor | 0.9 | Wang et al. (2016) |
| *M. edulis* | shell formation | 12528 | COL6A, collagen, type VI<br>Adhesome molecules | 0.9 | Dyachuk (2018) |
| *M. edulis* | shell formation | 8193 | mucin-13-like<br>Molluscan calcification | 0.9 | Marin et al. (2000) |
| *M. edulis* | Stress | 6739 | serine-protein kinase ATM<br>DNA damage sensor | 0.8 | Matsuoka and Igisu (2001) |

**Table 7.** *Cont.*

| Species | Category | ClusterID | Gene Names / Function (Ortholog cluster) | Ka/Ks | Ortholog reference |
|---|---|---|---|---|---|
| *M. galloprovincialis* | Immunity | 9813 | NIT1 | 2.5 | Lin et al. (2018) |
| *M. coruscus* | Shell formation | 8395 | Hemicentin (HMCN)<br>Extracellular ion-binding proteins in the biomineral matrix | 2 | Luo et al. (2015) |
| *M. coruscus* | Stress | 2491 | O-mannosyltransferase (TMTC)<br>Ca2+regulation and protein folding | 2 | Larsen et al. (2017) |
| *M. coruscus* | Immunity | 5530 | ADAR, adenosine deaminase<br>DNA binding. antiviral effectors | 1.6 | Green et al. (2015) |
| *M. coruscus* | Immunity | 9626 | Caveolin-1,Caveolin-3<br>Regulating neutrophil functional responses that underpin innate immunity | 1.2 | Zemans and Downey (2008) |
| *M. coruscus* | Immunity | 8044 | Apoptosis regulator BAX<br>Apoptosis regulator | 1.1 | Leprêtre et al. (2020) |
| *M. coruscus* | Stress | 17439 | 2-hydroxyglutaryl-CoA dehydratase (hgdC)<br>Iron-sulfur cluster binding | 1.1 | Locher et al. (2001) |
| *M. coruscus* | Immunity | 302 | Mannose receptor, C type (MRC)<br>Pathogen recognition receptor | 1 | Chen et al. (2015) |
| *M. coruscus* | Immunity | 6467 | Inhibitor of growth protein 1 (ING1)<br>Tumor suppressor gene | 1 | Garkavtsev et al. (1998) |
| *M. coruscus* | Immunity | 2098 | BIRC2_3<br>Physiological role in growth, immunity, and apoptosis | 1 | Wilson et al. (2016) |
| *M. coruscus* | Immunity | 8921 | Filamin<br>Recognition of pathogens | 1 | Maldonado-Aguayo et al. (2015) |
| *M. coruscus* | Immunity | 13594 | Proteasome regulatory (PSMD7)<br>Recognition of polyubiquitin chains and cleavage of ubiquitin from degraded proteins | 0.9 | Smits et al. (2020) |
| *M. coruscus* | Shell formation | 4996 | NOTCH1<br>Calcium signalling pathway and shell pigmentation | 0.9 | Auffret et al. (2020) |
| *M. coruscus* | Stress | 3709 | Heat shock protein 90kDa beta (HSP90B)<br>Heat shock protein | 0.9 | Cao et al. (2018) |
| *M. coruscus* | Immunity | 8284 | Cell division control protein 42<br>Roles in host defense | 0.9 | Xu et al. (2017) |
| *M. coruscus* | Stress | 5847 | Ammonium transporter (amt)<br>Ammonium transporter | 0.9 | Bu et al. (2019) |
| *M. coruscus* | Immunity | 12429 | RING finger protein 145<br>Ubiquitination | 0.8 | Cook et al. (2017) |
| *M. coruscus* | Stress | 12587 | Lysine methyltransferase 4 (EEF1AKMT4)<br>Enhances the function of heat shock factor 1 during the heat shock response | 0.8 | Vera et al. (2014) |
